# The unconventional cytoplasmic sensing mechanism for ethanol chemotaxis in *Bacillus subtilis*

**DOI:** 10.1101/2020.06.04.135640

**Authors:** Payman Tohidifar, Girija A. Bodhankar, Sichong Pei, C. Keith Cassidy, Hanna E. Walukiewicz, George W. Ordal, Phillip J. Stansfeld, Christopher V. Rao

## Abstract

Motile bacteria sense chemical gradients using chemoreceptors, which consist of distinct sensing and signaling domains. The general model is that the sensing domain binds the chemical and the signaling domain induces the tactic response. Here, we investigated the unconventional sensing mechanism for ethanol taxis in *Bacillus subtilis*. Ethanol and other short-chain alcohols are attractants for *B. subtilis*. Two chemoreceptors, McpB and HemAT, sense these alcohols. In the case of McpB, the signaling domain directly binds ethanol. We were further able to identify a single amino-acid residue Ala^431^ on the cytoplasmic signaling domain of McpB, that when mutated to a serine, reduces taxis to ethanol. Molecular dynamics simulations suggest ethanol binds McpB near residue Ala^431^ and mutation of this residue to serine increases coiled-coil packing within the signaling domain, thereby reducing the ability of ethanol to bind between the helices of the signaling domain. In the case of HemAT, the myoglobin-like sensing domain binds ethanol, likely between the helices encapsulating the heme group. Aside from being sensed by an unconventional mechanism, ethanol also differs from many other chemoattractants because it is not metabolized by *B. subtilis* and is toxic. We propose that *B. subtilis* uses ethanol and other short-chain alcohols to locate prey, namely alcohol-producing microorganisms.

**Importance:** Ethanol is a chemoattractant for *Bacillus subtilis* even though it is not metabolized and inhibits growth. *B. subtilis* likely uses ethanol to find ethanol-fermenting microorganisms for prey. Two chemoreceptors sense ethanol: HemAT and McpB. HemAT’s myoglobin-like sensing domain directly binds ethanol, but the heme group is not involved. McpB is a transmembrane receptor consisting of an extracellular sensing domain and a cytoplasmic signaling domain. While most attractants bind the extracellular sensing domain, we found that ethanol directly binds between inter-monomer helices of the cytoplasmic signaling domain of McpB, using a mechanism akin to those identified in many mammalian ethanol-binding proteins. Our results indicate that the sensory repertoire of chemoreceptors extends beyond the sensing domain and can directly involve the signaling domain.

## Introduction

Many bacteria move in response to external chemical gradients through a process known as chemotaxis (1). Typically, bacteria migrate up gradients of chemicals that support their growth and down ones that inhibit it. These chemicals are commonly sensed using transmembrane chemoreceptors, which consist of an extracellular sensing domain and a cytoplasmic signaling domain along with a cytoplasmic HAMP domain that couples the two domains. While a number of sensing mechanisms exist, the best understood one involves direct binding of the chemical to the extracellular sensing domain (2). In flagellated bacteria such as *Bacillus subtilis* and *Escherichia coli*, this binding event induces a conformational change in the cytoplasmic signaling domain that alters the autophosphorylation rate of an associated histidine kinase known as CheA (3). The phosphoryl group is then transferred to a soluble response regulator known as CheY, which modulates the swimming behavior of the bacterium by changing the direction of flagellar rotation. The chemical gradients themselves are sensed using a temporal mechanism involving sensory adaptation (4).

While many chemicals are sensed by the extracellular sensing domain, some are sensed by the cytoplasmic domains, typically using an indirect mechanism. For example, sugars transported by the phosphoenolpyruvate transfer system (PTS) are indirectly sensed through interactions between the PTS proteins and chemoreceptor signaling complexes (5, 6). In the case of *E. coli*, changes in intracellular pH are sensed by the cytoplasmic HAMP domain (7). In addition, changes in osmolarity are sensed through alterations in the packing of the chemoreceptors cytoplasmic signaling domains (8). To our knowledge, however, there have been no reports of direct sensing by the chemoreceptor cytoplasmic signaling domain. This has not been particularly surprising given that the cytoplasmic signaling domain, which consists of a long dimeric four-helix coiled-coil (9), lacks an obvious ligand-binding pocket.

In this work, we investigated chemotaxis to ethanol in *B. subtilis*. This short-chain alcohol is an attractant for *B. subtilis* even though it is not used as a carbon source and inhibits cell growth. Ethanol is directly sensed by two chemoreceptors, HemAT and McpB. Sensing by HemAT fits the conventional model where ethanol binds the sensing domain. However, in the case of McpB, we found that ethanol is directly sensed by the cytoplasmic signaling domain using a mechanism analogous to many eukaryotic ethanol-binding proteins.

## Results

### *B. subtilis* exhibits chemotaxis to short-chain alcohols

We employed the capillary assay to measure *B. subtilis* chemotaxis to alcohols with increasing chain lengths (C1 to C5). The resulting data show that *B. subtilis* exhibits chemotaxis to methanol, ethanol, 2-propanol, and *tert*-butanol. No significant responses to 1-propanol, 1-butanol, and 1-pentanol were observed (**Fig. 1A**). To elucidate the underlying sensing mechanism, we focused on ethanol, because it is produced and utilized by a wide range of microorganisms in nature (10). We first measured the response to increasing ethanol concentration using the capillary assay. (**Fig. 1B**). Unlike many other attractants such as amino acids (11–13), a tactic response to ethanol was only observed at relatively high concentrations (> 50 mM versus 1-100 μM for amino acids). The ethanol response peaked at 1.78 M (∼ 10% (v/v)). The response decreased at higher concentrations, most likely due to ethanol being toxic at these concentrations (14).

**FIG 1.**
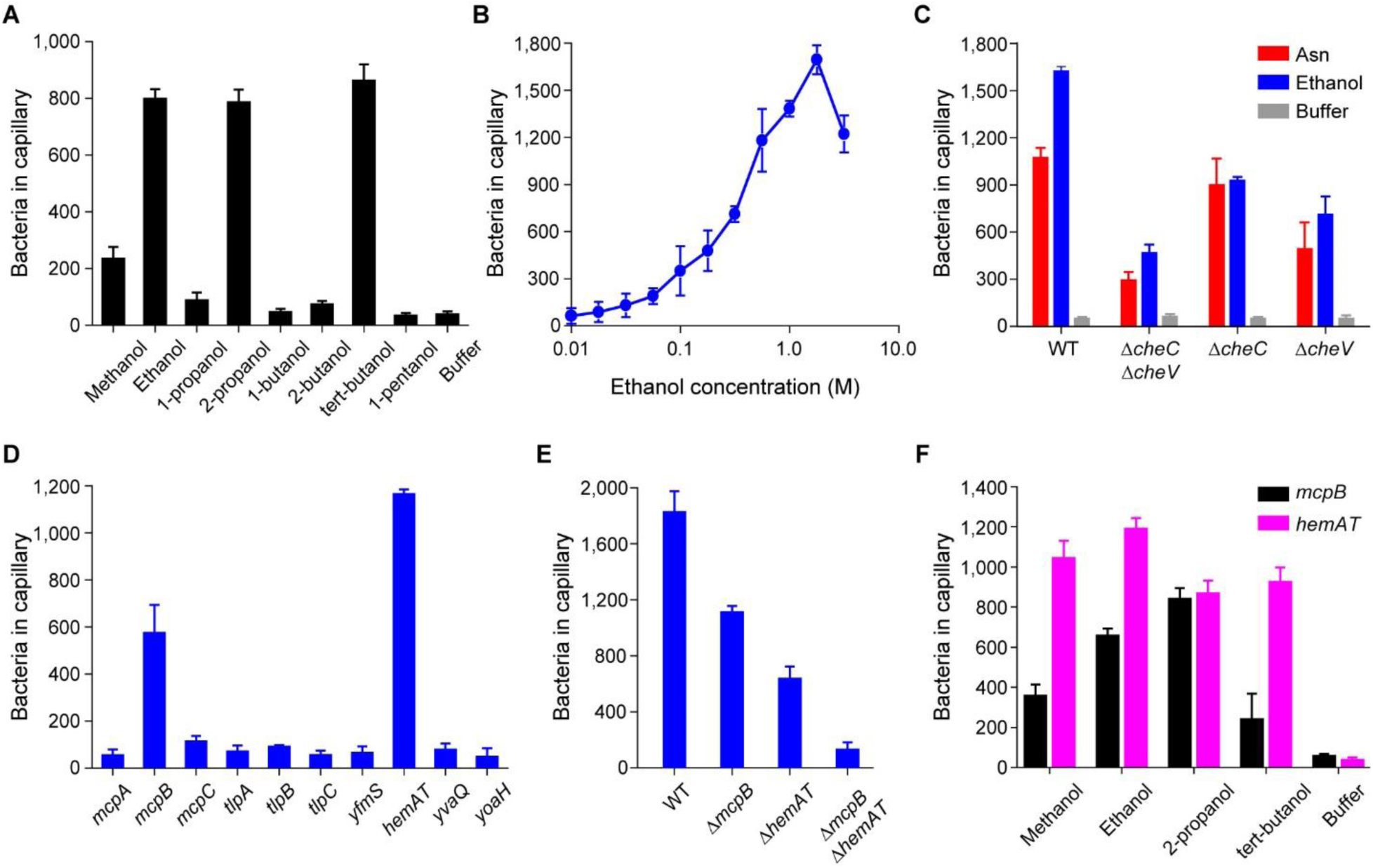
*B. subtilis* exhibits chemotaxis toward short-chain alcohols. (A) Responses of the wild-type strain to 0.5 M short-chain alcohols with increasing chain lengths (C1 to C5). (B) Dose-dependent response of the wild-type strain to increasing concentration of ethanol. (C) Responses of adaptation-deficient mutants to ethanol and asparagine. (D) Responses of mutants expressing single chemoreceptors to ethanol. (E) Responses of mutants lacking key chemoreceptors to ethanol. (F) Responses of mutants expressing McpB or HemAT as their sole chemoreceptor to short-chain alcohols. In these experiments, ethanol and asparagine concentrations were 1.78 M and 3.16 µM, respectively, unless otherwise mentioned. Negative control responses of the strains expressing single chemoreceptor to buffer were all under 100 colonies per capillary. Error bars denote the standard deviations from three biological replicates performed on three separate days.

### All three adaptation systems contribute to ethanol taxis

*B. subtilis* employs three adaptation systems – the methylation, CheC/CheD/CheYp, and CheV systems – for sensing chemical gradients (4, 15). To test whether these adaptation systems are involved in ethanol taxis, we employed mutants where these systems were selectively inactivated. We first tested ethanol taxis using a mutant (Δ*cheC* Δ*cheV*) where the CheC/CheD/CheYp and CheV adaptation systems were inactivated, leaving only the methylation system functional. Taxis to both ethanol and asparagine, which was used as a control, was reduced 30% in this mutant (**Fig. 1C**). We also observed reduced taxis in the Δ*cheC* and Δ*cheV* mutants, though the reduction was less than what was observed with the double mutant. Interestingly, CheC/CheD/CheYp system appears to be more important for sensing ethanol gradients than for asparagine gradients (**Fig. 1C**). We did not test a Δ*cheR* Δ*cheB* mutant, which lacks the two enzymes involved in methylation system, because this mutant exhibits poor motility in general due to excessive tumbling.

### McpB and HemAT are the chemoreceptors for short-chain alcohols

*B. subtilis* has ten chemoreceptors (16). To determine the chemoreceptors involved in ethanol taxis, we first tested mutants expressing just one chemoreceptor using the capillary assay. Only strains expressing McpB or HemAT as their sole chemoreceptor were capable of ethanol taxis (**Fig. 1D**). The response was greater for strains expressing HemAT, suggesting that it is the main receptor for ethanol taxis. This is not surprising as HemAT is more highly expressed than McpB (19,000 versus 6,200) (16). We next tested the effect of deleting these chemoreceptors in the wild type. When either McpB or HemAT was deleted (Δ*mcpB* or Δ*hemAT*), we observed reduced taxis toward ethanol. The reduction was greater in the Δ*hemAT* mutant, again suggesting that HemAT is the main receptor for ethanol taxis. When both chemoreceptors were deleted in the wild type (Δ*mcpB* Δ*hemAT*), ethanol taxis was almost completely eliminated (**Fig. 1E**). We also found that strains expressing McpB or HemAT as their sole chemoreceptor responded to methanol, 2-propanol, and *tert*-butanol (**Fig. 1F**). Strains expressing HemAT as their sole chemoreceptor exhibited stronger responses to these alcohols than strains expressing McpB alone with the exception of 2-propanol where the responses were similar.

### Chemotaxis to ethanol is independent of its metabolism

Many bacteria metabolize ethanol (17). One possibility is that *B. subtilis* senses products of ethanol metabolism rather than ethanol itself. Indeed, such a mechanism occurs in *Pseudomonas putida* with regards to alcohol taxis (18). Therefore, we tested whether *B. subtilis* can grow on ethanol (**Fig. 2A**). These growth experiments were performed using the parental strain *B. subtilis* 168, which lacks the auxotrophies present in the chemotaxis strain OI1085. When cells were cultured in minimal medium with ethanol as the sole carbon source, no growth was observed. However, the cells did grow when ethanol was replaced with glucose. We also tested *B. subtilis* 168 growth in rich medium containing different amounts of ethanol to determine whether the cells were able to consume ethanol even though ethanol alone does not support growth as the sole carbon source. While the cells were able to grow in rich medium containing ethanol, no decreases in ethanol concentrations were observed (**Fig. 2B**). These results indicate that *B. subtilis* does not consume ethanol.

**FIG 2.**
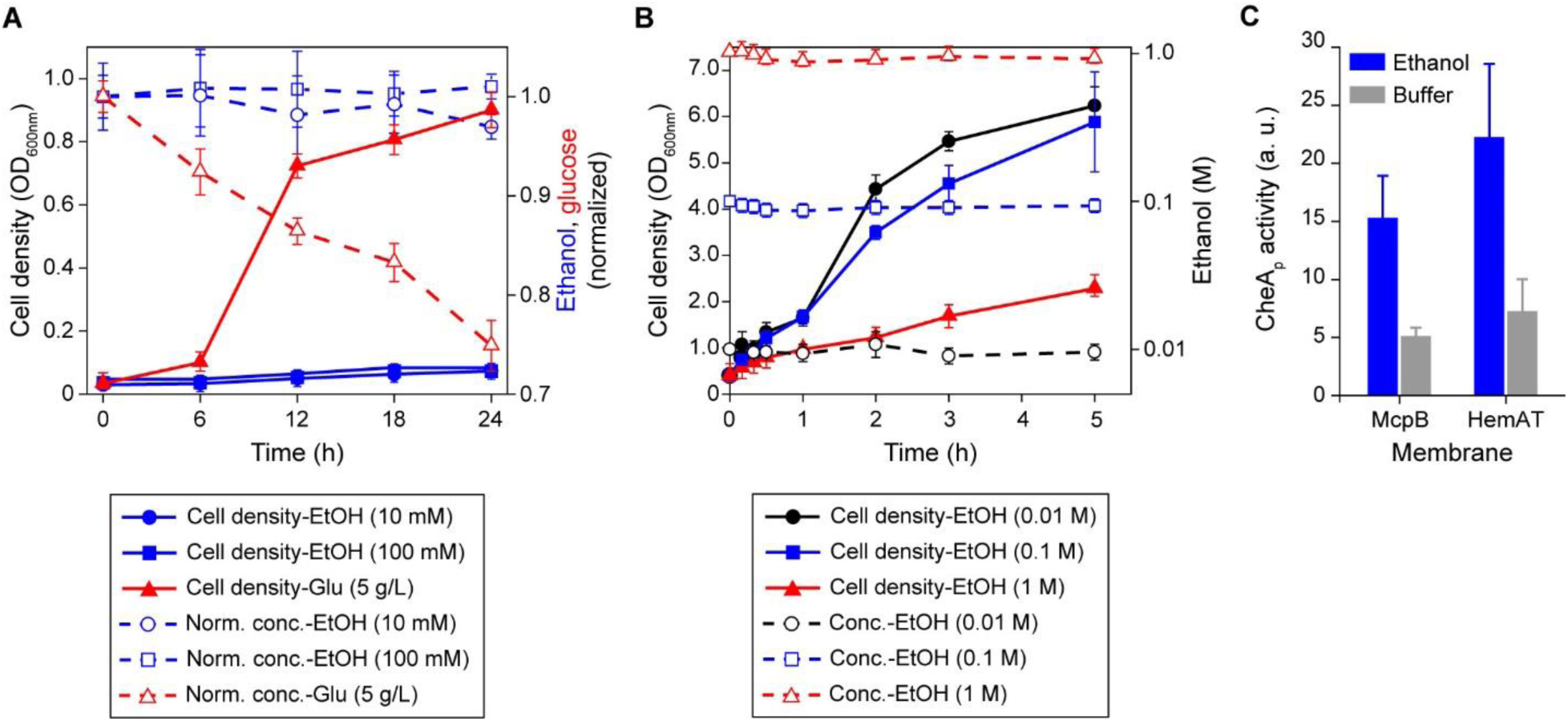
*B. subtilis* chemotaxis to ethanol is independent of its metabolism. (A) Cell growth in minimal medium supplemented with 10 mM ethanol (blue solid circles), 100 mM ethanol (blue solid squares), and 5 g/L glucose (red solid triangles) tested as a positive control. Dashed lines with the corresponding symbols depict normalized concentrations of chemicals measured over the course of 24 h. (B) Cell growth in rich medium containing 10 mM ethanol (black sold circles), 100 mM ethanol (blue solid squares), and 1 M ethanol (red solid triangles). Dashed lines with open symbols depict absolute concentration of ethanol measured at three different conditions over the course of 5 h. (C) Levels of phosphorylated CheA kinase complexed with CheW, CheD, and McpB or HemAT within the isolated membranes, in presence of 1 M ethanol or buffer, as negative control. Error bars denote the standard deviations from three biological replicates performed on three separate days.

Oxidation of alcohols to acetaldehyde and subsequently to carboxylic acids can potentially change the redox state of the cells. This change could possibly be perceived as a sensory signal through a process known as energy taxis (19). *B. subtilis* can ferment glucose to acetate and ethanol when grown in presence of pyruvate or a mixture of amino acids (20, 21). In this process, alcohol dehydrogenase (ADH) reduces acetaldehyde to ethanol using NADH as the cofactor. Whether ADH can oxidize ethanol to acetaldehyde in *B. subtilis* is unknown. To test whether this occurs, we measured ADH activities using *B. subtilis* cell lysates prepared from aerobic and anaerobic cultures. As a positive control, ADH activities using *E. coli* cell lysates were also measured (22). No ADH activity was observed with *B. subtilis* lysates whereas *E. coli* lysates obtained from anaerobic cultures had an ADH activity of 31.25 ± 1.85 units/mL. As expected, no ADH activity was detected with aerobic *E. coli* lysates. These results suggest that ethanol taxis in *B. subtilis* is independent of ethanol catabolism and is instead sensed directly by McpB and HemAT.

### Ethanol induces receptor-coupled kinase activity

We next performed an *in vitro* receptor-coupled kinase assay to test whether ethanol is able to activate CheA kinase (23). This assay has been used to study how attractant binding to chemoreceptors modulates CheA kinase activity (15, 23). Briefly, membranes expressing either McpB or HemAT were isolated. The chemotaxis signaling proteins CheA, CheW, and CheD were then added to these membranes to final concentrations that matched their stoichiometry in wild-type cells. Using this assay, we found that ethanol activates CheA kinase in a dose-dependent manner with membranes containing either McpB or HemAT as the sole chemoreceptor. Ethanol concentrations as low as 10 mM were sufficient to activate CheA kinase in both cases (**Fig. S1**). Additionally, kinase activation was stronger with HemAT-containing membranes (**Fig. 2C** and **Fig. S1**), which is consistent with our *in vivo* results. These results indicate that ethanol can induce chemotaxis signaling *in vitro*. This assay, however, is unable to determine whether ethanol directly interacts with the chemoreceptors, because the membranes might contain associated proteins that could be involved in signaling.

### McpB cytoplasmic signaling domain is involved in ethanol sensing

We next investigated ethanol taxis using receptor chimeras involving McpB to provide further insight regarding the sensing mechanism (24–26). We focused on McpB due to its high amino-acid similarity (57% to 65%) with three other *B. subtilis* chemoreceptors: McpA, TlpA, and TlpB. These four chemoreceptors all employ the same double Cache 1 domain for their sensing domain (2) and a highly conserved coiled-coil structure for their cytoplasmic signaling domain (9) (**Fig. 3A** and **3B**). Unlike McpB, HemAT is not a transmembrane chemoreceptor. We attempted to construct chimeras involving HemAT, McpA, and YfmS, another soluble chemoreceptor. However, none were functional in the sense that they did not respond to ethanol or molecular oxygen, which is the conventional attractant for HemAT (27).

**FIG 3.**
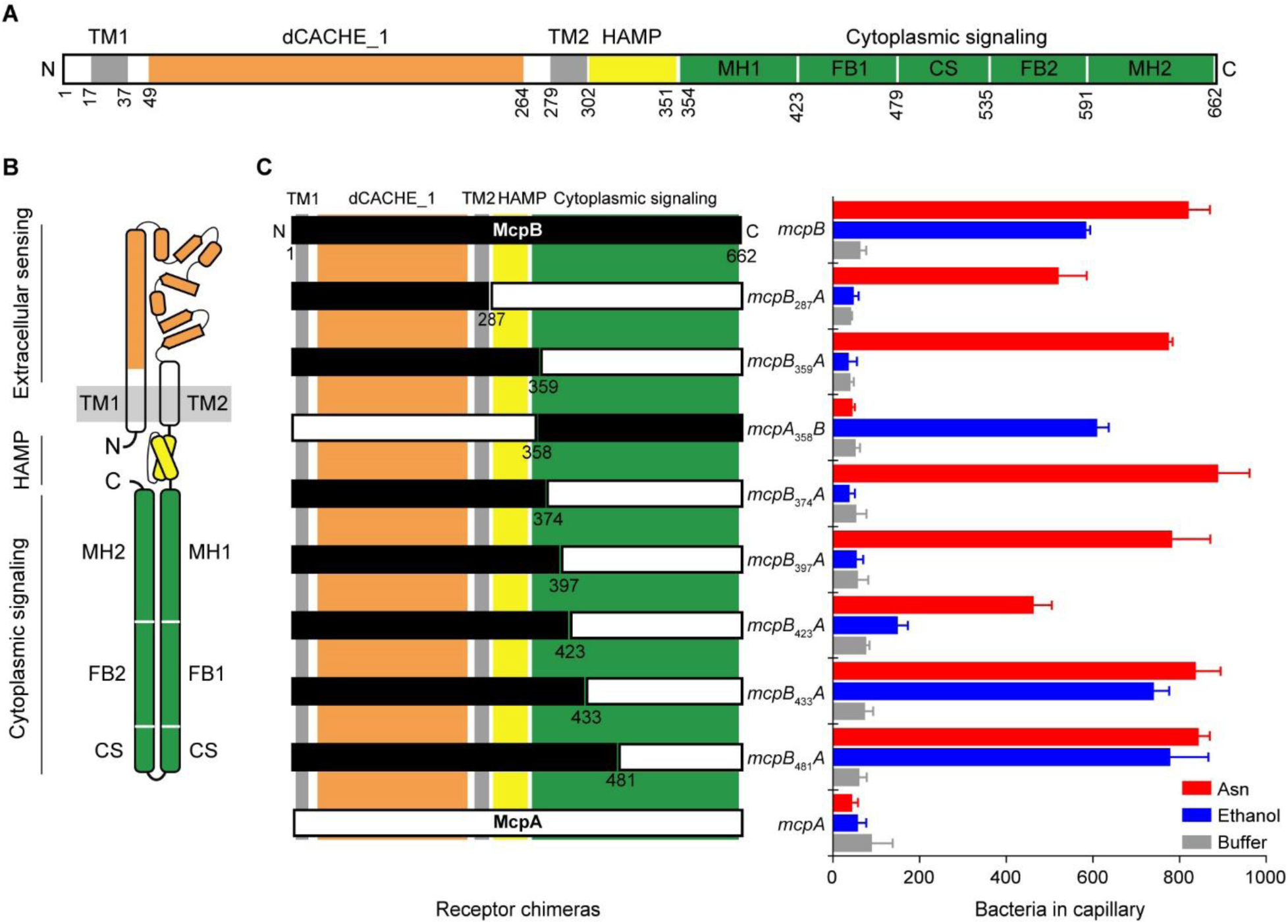
McpB cytoplasmic signaling domain is involved in ethanol sensing. (A) Domain structure of McpB, McpA, TlpA, and TlpB. All four chemoreceptors consist of an extracellular sensing domain with dCACHE_1 structure (orange) followed by transmembrane, TM1 and TM2 (gray), HAMP (yellow), and cytoplasmic signaling (green) domains. Three subdomains of the cytoplasmic signaling domain classified as methylation (adaptation) helices (MH), flexible (coupling) bundle (FB), and conserved signaling (protein contact region) (CS) tip are shown. (B) Cartoon structure of a monomer of the chemoreceptors. (C) Responses of mutants expressing chimeric receptors between McpA (white) and McpB (black) to 1.78 M ethanol, 3.16 µM asparagine, and buffer. Error bars denote the standard deviations from three biological replicates performed on three separate days.

We created chimeras between McpB and McpA, because the latter is not involved in ethanol taxis. In addition, we also measured the response to asparagine, because it is a known attractant for McpB, but not McpA, and binds the extracellular sensing domain (11). We first fused the N-terminal region of McpB to the C-terminal region of McpA: *mcpB*_287_*A* and *mcpB*_359_*A*. We then tested whether strains expressing these chimeras as their sole chemoreceptor respond to ethanol using the capillary assay. Both mutants did not respond to ethanol even though they still responded to asparagine (**Fig. 3C**). These results demonstrate that the extracellular sensing and cytoplasmic HAMP domains are not involved in sensing ethanol. Rather, the cytoplasmic signaling domain is involved. To verify our hypothesis, we tested a *mcpA*_358_*B* chimera. As expected, strain expressing *mcpA*_358_*B* as its sole chemoreceptor responded to ethanol. This strain, however, does not respond to asparagine, because it lacks the requisite McpB sensing domain (**Fig. 3C**).

A key feature of chemoreceptor cytoplasmic signaling domains are the characteristic heptad repeats (labeled *a* to *g*) associated with their coiled-coil structure, where each repeat is equivalent to two helical turns. Based on sequence conservation and structural analysis of heptads from several bacterial and archaeal chemoreceptors, the cytoplasmic signaling domains are classified into three structurally distinct subdomains. These subdomains are known as the methylation (adaptation) helices, the flexible (coupling) bundle, and the conserved signaling tip (protein contact region) (9) (**Fig. 3** and **Fig. S2**). To narrow down the region on these subdomains involved in ethanol sensing, we created *mcpB*_374_*A*, *mcpB*_397_*A*, *mcpB*_423_*A, mcpB*_433_*A,* and *mcpB*_481_*A* chimeras. Strains expressing *mcpB*_374_*A* and *mcpB*_397_*A* as their sole chemoreceptor did not respond to ethanol even though they still responded to asparagine. Strains expressing *mcpB*_423_*A* as their sole chemoreceptor exhibited a reduced response to ethanol and asparagine. However, when *mcpB*_433_*A* and *mcpB*_481_*A* were tested, the corresponding chimera expressing strains were able to respond to ethanol and asparagine at levels similar to the wild-type control (**Fig. 3C**). These results suggest that the region spanning the residues 397 to 433 on McpB is involved in sensing ethanol. Furthermore, the region spanning the residues 423 to 433 on McpB appears to be the principal region involved in ethanol sensing.

### McpB residue involved in ethanol sensing

The region spanning residues 397 to 433 on McpB is necessary for ethanol taxis. As a first step toward identifying the binding site, we performed *in silico* docking experiments with ethanol and the McpB dimer fragment spanning residues 390 to 435 on the N-helix and neighboring residues 577 to 622 on the C-helix. The resulting data from the docking analysis yielded five distinct clusters of putative amino acid residues involving both the N-helix and the C-helix of the dimer fragment (**Table S1** and **Fig. S3A**). We next aligned the amino-acid sequences spanning residues 392 to 434 on the N-helix and neighboring residues 578 to 620 on the C-helix of McpB, McpA, TlpA, and TlpB. (**Fig. S3B**). Among the 20 putative binding residues, Thr^424^, Asp^427^, and Ala^431^ on the N-helix and Glu^581^ and Lys^585^ on the C-helix were not conserved between the four chemoreceptors and, thus, were targeted for mutational analysis (**Fig. 4A** and **4B**). Mutants expressing *mcpB-T424A, mcpB-D427T, mcpB-E581Q,* and *mcpB-K585E* as their sole chemoreceptor exhibited responses to ethanol similar to the wild-type *mcpB*. However, the strain expressing *mcpB-A431S* as its sole chemoreceptor failed to respond to ethanol. In addition, all strains supported asparagine taxis, indicating that these mutated receptors were functional (**Fig. 4C**). We also measured the response of the strain expressing *mcpB-A431S* to methanol, 2-proponal, and *tert*-butanol in the capillary assay and observed reduced responses (**Fig. 4D**), suggesting that Ala^431^ is an important residue for alcohol taxis overall.

**FIG 4.**
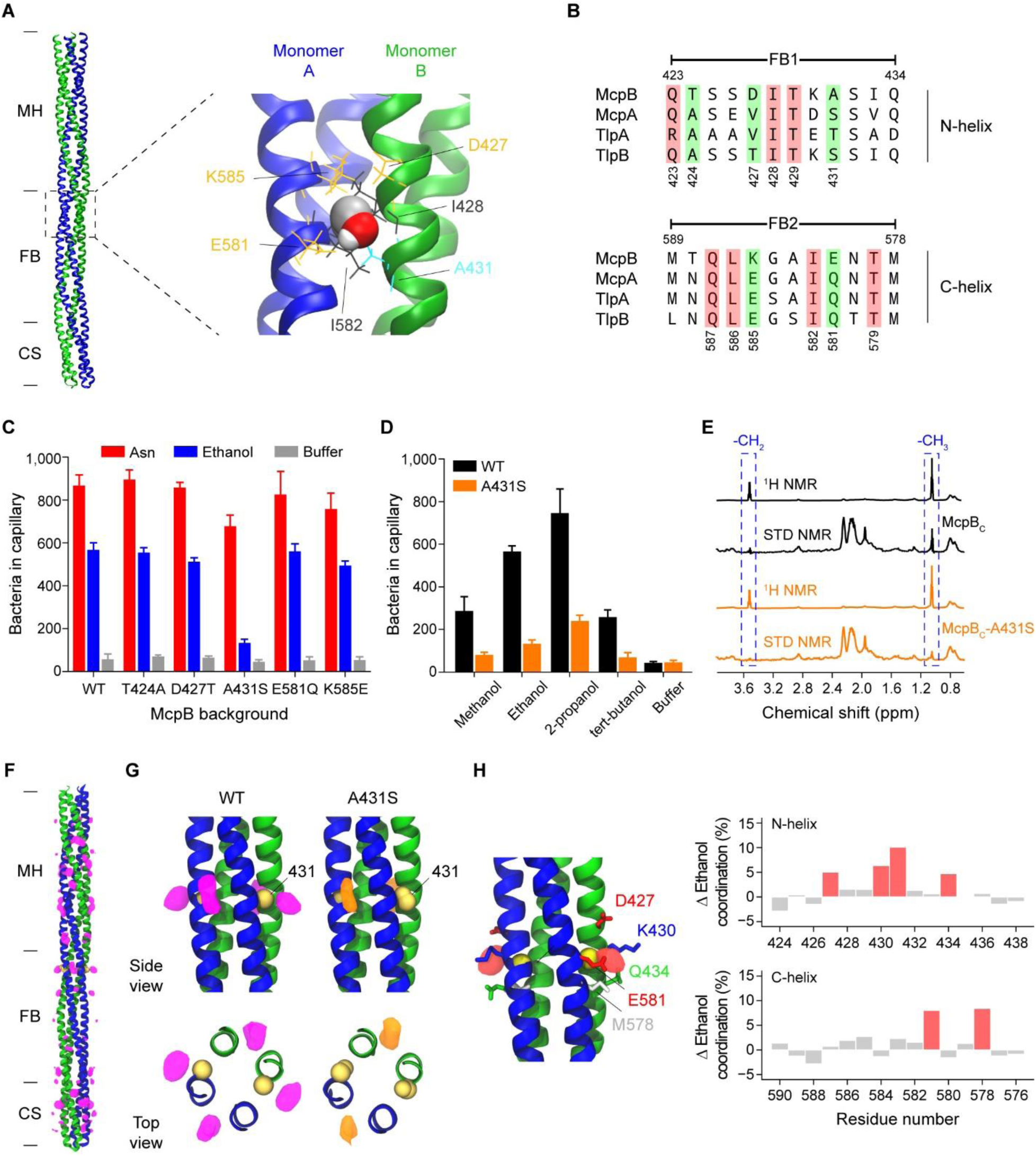
Alcohols are directly sensed by the cytoplasmic signaling domain of McpB (A) A putative binding site within the primary ethanol-sensing region spanning residues (423 to 433) on the N-helix and the neighboring residues (579 to 589) on the C-helix of the McpB cytoplasmic signaling domain. (B) Amino acid sequence alignment of the primary ethanol-sensing region for McpB and the corresponding regions on McpA, TlpA, and TlpB. (C) Responses of strains expressing McpB mutants as their sole chemoreceptors to 1.78 M ethanol, 3.16 µM asparagine, and buffer. (D) Responses of strains expressing wild-type McpB and the McpB-A431S mutant as their sole chemoreceptor to 1.78 M short-chain alcohols and buffer. (E) ^1^H and STD-NMR spectra for 50 µM wild-type and mutant (A431S) recombinant McpB cytoplasmic region (McpB_C_) spanning residues (305 to 662). Two peaks at 1.05 ppm and 3.51 ppm (shown inside dashed boxes) respectively correspond to -CH_3_ and -CH_2_ epitopes of ethanol. (F) Density map of the average ethanol occupancy (purple) along the wild-type McpB cytoplasmic signaling domain (McpB_C_) spanning residues 352 to 662, as predicted by MD simulation. (G) Enlarged side and top views of the ethanol occupancy surrounding the residue 431 (yellow) in the wild-type (purple) and the A431S mutant (orange) McpB_C_. (H) Difference map (red density) between the wild-type and the A431S mutant McpB_C_ surrounding the residue 431, highlighting the loss of an inter-monomer ethanol binding site in the A431S mutant. Changes in protein-ethanol coordination highlight the putative amino-acid residues (red bars) involved in ethanol binding. Error bars reported in panels C and D denote the standard deviations from three biological replicates performed on three separate days. Ethanol occupancy and coordination values are generated from three independent MD simulations.

**TABLE 1.**
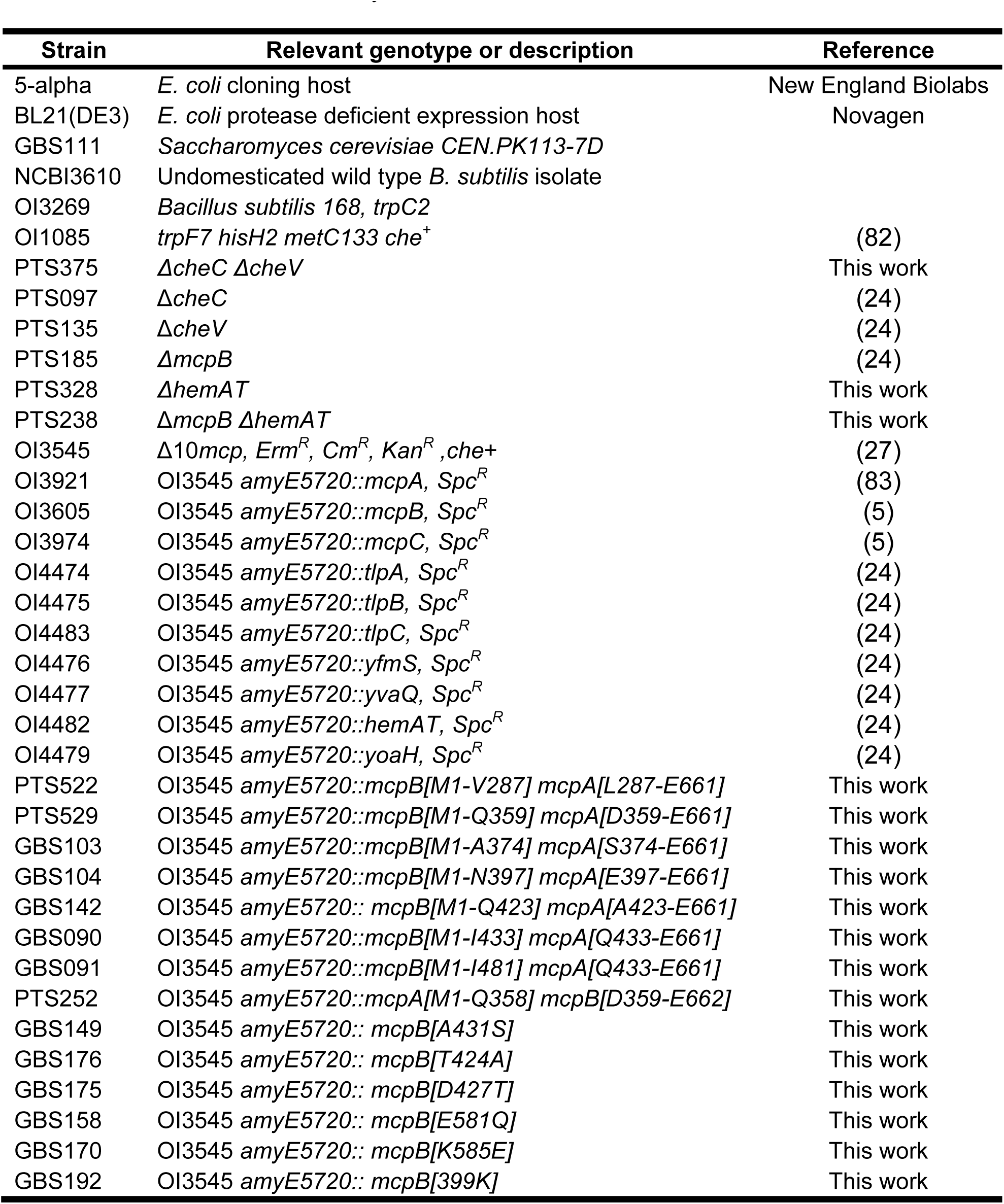
Strains used in this study.

| Strain | Relevant genotype or description | Reference |
| --- | --- | --- |
| 5-alpha | *E. coli* cloning host | New England Biolabs |
| BL21(DE3) | *E. coli* protease deficient expression host | Novagen |
| GBS111 | *Saccharomyces cerevisiae* CEN.PK113-7D |  |
| NCBI3610 | Undomesticated wild type *B. subtilis* isolate |  |
| OI3269 | *Bacillus subtilis* 168, *trpC2* |  |
| OI1085 | *trpF7 hisH2 metC133 che<sup>+</sup>* | (82) |
| PTS375 | $\Delta cheC \Delta cheV$ | This work |
| PTS097 | $\Delta cheC$ | (24) |
| PTS135 | $\Delta cheV$ | (24) |
| PTS185 | $\Delta mcpB$ | (24) |
| PTS328 | $\Delta hemAT$ | This work |
| PTS238 | $\Delta mcpB \Delta hemAT$ | This work |
| OI3545 | $\Delta 10mcp, Erm^R, Cm^R, Kan^R, che^+$ | (27) |
| OI3921 | OI3545 *amyE5720::mcpA, Spc<sup>R</sup>* | (83) |
| OI3605 | OI3545 *amyE5720::mcpB, Spc<sup>R</sup>* | (5) |
| OI3974 | OI3545 *amyE5720::mcpC, Spc<sup>R</sup>* | (5) |
| OI4474 | OI3545 *amyE5720::tlpA, Spc<sup>R</sup>* | (24) |
| OI4475 | OI3545 *amyE5720::tlpB, Spc<sup>R</sup>* | (24) |
| OI4483 | OI3545 *amyE5720::tlpC, Spc<sup>R</sup>* | (24) |
| OI4476 | OI3545 *amyE5720::yfmS, Spc<sup>R</sup>* | (24) |
| OI4477 | OI3545 *amyE5720::yvaQ, Spc<sup>R</sup>* | (24) |
| OI4482 | OI3545 *amyE5720::hemAT, Spc<sup>R</sup>* | (24) |
| OI4479 | OI3545 *amyE5720::yoaH, Spc<sup>R</sup>* | (24) |
| PTS522 | OI3545 *amyE5720::mcpB[M1-V287] mcpA[L287-E661]* | This work |
| PTS529 | OI3545 *amyE5720::mcpB[M1-Q359] mcpA[D359-E661]* | This work |
| GBS103 | OI3545 *amyE5720::mcpB[M1-A374] mcpA[S374-E661]* | This work |
| GBS104 | OI3545 *amyE5720::mcpB[M1-N397] mcpA[E397-E661]* | This work |
| GBS142 | OI3545 *amyE5720::mcpB[M1-Q423] mcpA[A423-E661]* | This work |
| GBS090 | OI3545 *amyE5720::mcpB[M1-I433] mcpA[Q433-E661]* | This work |
| GBS091 | OI3545 *amyE5720::mcpB[M1-I481] mcpA[Q433-E661]* | This work |
| PTS252 | OI3545 *amyE5720::mcpA[M1-Q358] mcpB[D359-E662]* | This work |
| GBS149 | OI3545 *amyE5720::mcpB[A431S]* | This work |
| GBS176 | OI3545 *amyE5720::mcpB[T424A]* | This work |
| GBS175 | OI3545 *amyE5720::mcpB[D427T]* | This work |
| GBS158 | OI3545 *amyE5720::mcpB[E581Q]* | This work |
| GBS170 | OI3545 *amyE5720::mcpB[K585E]* | This work |
| GBS192 | OI3545 *amyE5720::mcpB[399K]* | This work |

### Ethanol directly binds to the McpB cytoplasmic signaling domain

To test whether ethanol directly interacts with McpB, we conducted saturation-transfer difference nuclear magnetic resonance (STD-NMR) experiments using recombinant McpB. STD-NMR has been used to measure weak interactions between proteins and their ligands (28–31). Briefly, in these experiments the protein is selectively saturated at specific frequencies. The magnetization is then transferred to the surrounding, low molecular-weight ligands in a distance-dependent manner. The ligand epitopes in close proximity of the protein receive higher saturation (32), implying direct binding to the protein.

We first tested the McpB cytoplasmic region (McpB_C_) spanning residues 305 to 662, which corresponds to the HAMP and the signaling domains (see Fig. 3A). The resulting ^1^H spectra for the McpB_C_ protein incubated with 3 mM ethanol (60-fold excess of the protein) is shown in **Fig. 4E**. Two peaks for ethanol appeared near 1.05 ppm and 3.51 ppm, which respectively correspond to -CH_3_ and -CH_2_ epitopes of ethanol. Ligand signals were also observed at the expected chemical shift values (1.05 ppm and 3.51 ppm) on the STD spectra. Additionally, the area under the STD peak corresponding to the -CH_2_ epitope was about five-fold (18%) less than that of the -CH_3_ epitope (**Fig. 4E**), suggesting that the -CH_3_ moiety of ethanol is closer to the protein than its -CH_2_ moiety. These results collectively indicate that ethanol directly interacts with the McpB cytoplasmic region.

Strains expressing *mcpB-*A431S as their sole chemoreceptor exhibited a reduced response to ethanol when tested in the capillary assay (**Fig. 4C**). To determine whether the A431S mutation reduces ethanol binding, we repeated the STD-NMR experiments with recombinant McpB_C_-A431S protein. Because single mutations may impair proper folding of proteins, we first measured the circular dichroism spectra for both the wild-type McpB_C_ and the McpB_C_-A431S proteins. We observed similar spectra for both proteins, which suggests that the mutant protein was properly folded (**Fig. S4**). We then performed STD-NMR experiments with the McpB_C_-A431S in presence of 3 mM ethanol.

The resulting STD spectra showed reduced peaks near 1.05 ppm and 3.51 ppm as compared to wild-type McpB_C_ (**Fig. 4E**). The saturation fraction of ethanol, which corresponds to the ratio of areas under the respective -CH_3_ peaks on STD and ^1^H spectra, is 0.23 for McpB_C_ and 0.1 for the McpB_C_-A431S. These results imply that the residue Ala^431^ has a role in ethanol binding to the McpB signaling domain.

### Molecular dynamics simulation suggests ethanol directly binds the residue Ala^431^

To gain insight regarding the ethanol binding mechanism, we performed molecular dynamics (MD) simulations of the wild-type and A431S McpB cytoplasmic signaling dimers (residues 352 to 662) in presence of ethanol. Our simulations demonstrate that ethanol can bind nonspecifically throughout the cytoplasmic signalling domain in both the wild-type and the mutant McpB dimers, primarily interacting along the inter-helical grooves of the four-helix bundle (**Fig. 4F** and **Fig. S5A**). A comparison of the ethanol occupancy between the wild-type and A431S mutant McpB shows little variation overall but exhibits a marked difference in the region immediately surrounding residue 431. In particular, while ethanol was observed to bind at both the inter- and intra-monomer interfaces in the wild-type simulations, the inter-monomer binding site associated with the residue 431 side chain was not present in the mutant simulations (**Fig. 4G**), suggesting that the A431S mutation reduces the binding ability of ethanol at residue 431. Indeed, within the flexible-bundle region, the residues displaying the greatest change in ethanol coordination between the wild-type and the A431S mutant form a concentric pocket centered on residue 431 at the inter-monomer interface (**Fig. 4H**).

Our analyses identified another interesting feature of ethanol binding, namely that it is able to penetrate the surface of the McpB cytoplasmic domain to bind within the core of the coiled coil. In particular, we observed that ethanol entered between the individual helices of the four-helix bundle at two locations in the methylation-helix region: one involving N-helix residues 393-400 and C-helix residues 613-617 and another involving N-helix residues 382-387 and C-helix residues 628-631 (**Fig. S5A**). While ethanol binding to these regions was observed in both the wild-type and A431S mutant simulations, the wild-type binding events resulted in longer dwell times, giving rise to the difference in ethanol coordination observed in these regions (**Fig. S5A**). Preliminary analysis of the two sites, however, suggests they do not themselves play a significant role in signaling. The latter is located outside the region involved in ethanol sensing (see **Fig. 3C**) and the former, except for residue Glu^399^, is highly conserved among the four chemoreceptors (see Fig. S3B). Indeed, we did not observe a significant reduction in response to ethanol compared to the wild-type control when we tested a mutant expressing *mcpB-E399K* as its sole chemoreceptor in the capillary assay (569 ± 29.1 cells versus 586.1 ± 9.0 cells, respectively). Nevertheless, these observations hint at a signaling mechanism in which ethanol may penetrate to the core of the cytoplasmic domain where it can affect the packing and overall stability of the bundle.

To further investigate the above packing hypothesis, we analyzed the strength of knobs-in-holes interactions in the region surrounding residue 431 over the course of the simulations. We observed that the A431S mutation leads to increased occupancy of the 431 knob itself as well as nearby knobs on the C-helix at positions 583 and 585 (**Fig. S5B**), indicating stronger hydrophobic interactions between the individual helices.

Therefore, our simulation results suggest that the A431S mutation, which decreases the McpB ethanol response, not only reduces the direct binding of ethanol but also strengthens coiled-coil packing in the region. One possibility is that the reduced local concentration of ethanol and improved packing in the A431S mutant decreases the ability of ethanol to intercalate with the knobs-into-holes interactions near residue 431 and, thus, its ability to induce signaling.

### The HemAT sensing domain helices are involved in direct ethanol sensing

HemAT is a cytoplasmic chemoreceptor, which consists of an N-terminal sensing domain and a C-terminal signaling domain. To determine which domain is involved in ethanol sensing, we conducted the STD-NMR experiments with the purified sensing domain (HemAT_N_), spanning residues 1 to 178, and the purified signaling domain (HemAT_S_), spanning residues 177 to 432 of the HemAT, in presence of 3 mM ethanol.

The resulting ^1^H and STD spectra with the HemAT_N_ showed clear peaks near 1.05 ppm and 3.51 ppm, which correspond to the -CH_3_ and the -CH_2_ moieties of ethanol. Additionally, the ratio of areas in the STD spectra compared to ^1^H spectra was 0.27 for the -CH_3_ moiety and 0.85 for the -CH_2_ moiety, suggesting that -CH_2_ moiety of ethanol is closer to the protein than its -CH_3_ moiety. The STD spectra with the HemAT signaling domain, however, showed negligible peaks near the expected chemical shift values (1.05 ppm and 3.51 ppm) (**Fig. 5A**). These results collectively indicate that ethanol binds the sensing domain of the HemAT.

**FIG 5.**
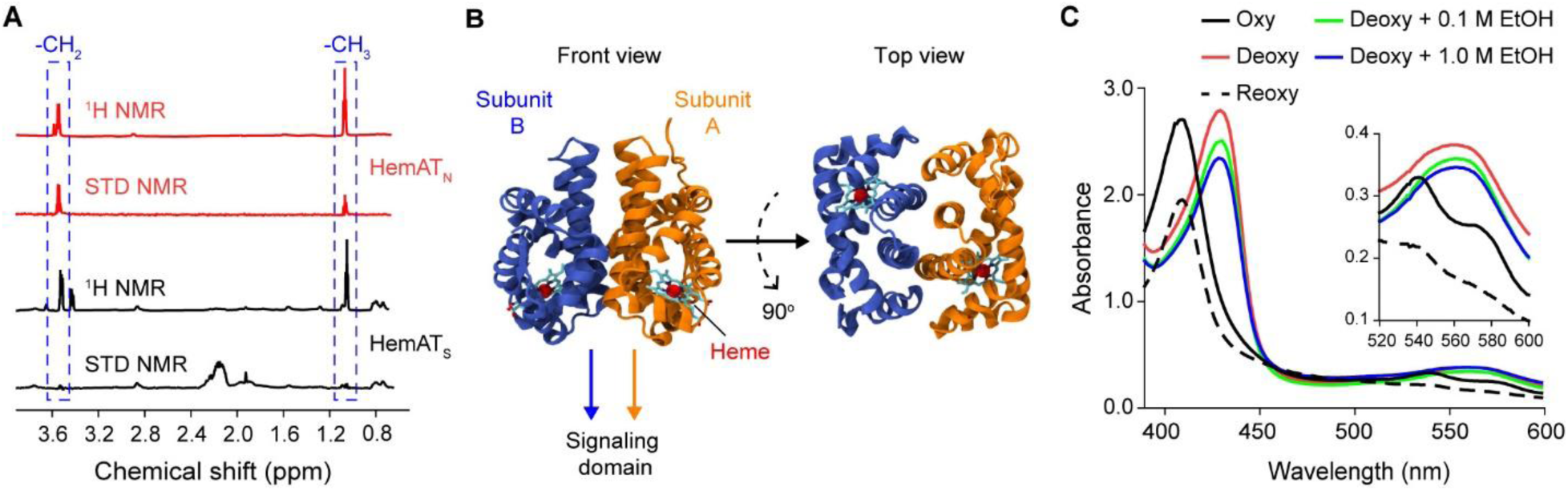
Ethanol directly binds within the helices of the HemAT sensing domain. (A) ^1^H and STD-NMR spectra for 50 µM recombinant HemAT C-terminal signaling domain (HemAT_S_) spanning residue (177 to 432) and for 50 µM recombinant HemAT N-terminal sensing domain (HemAT_N_) spanning residues (1 to 178), in presence of 3 mM ethanol. Two peaks at 1.05 ppm and 3.51 ppm (shown inside dashed boxes) respectively correspond to -CH_3_ and -CH_2_ epitopes of ethanol. (B) Crystal structure of the dimeric HemAT sensing domain. (C) UV-spectra of recombinant HemAT N-terminal sensing domain (HemAT_N_) in absence and presence of molecular oxygen, 0.1 M ethanol, and 1.0 M ethanol.

The sensing domain of the HemAT dimer is composed of a four-helical bundle as its core and a heme group in each subunit (**Fig. 5B**), which is known to bind molecular oxygen (33). UV-spectral analyses have shown that the oxygen molecule binds the heme group by forming hydrogen bonds with 6-coordinate ferrous heme (34, 35). To determine whether the heme group also interacts with ethanol, we conducted UV spectroscopy experiments with the purified HemAT sensing domain (HemAT_N_) and ethanol. As a control, we first measured UV absorption of both oxygenated and deoxygenated forms of the protein to verify that the heme group on the purified protein is functional. Consistent with the previous reports (27, 34, 36), the oxygenated form of the HemAT_N_ exhibited three major canonical peaks at 412 nm (Soret), 544 nm (β-band), and 578 nm (α-band), and the dithionite-reduced deoxygenated form of the protein exhibited two major peaks at 434 nm and 556 nm (**Fig. 5C**). Next, we measured UV absorption of the deoxygenated HemAT sensing domain in presence of varying concentrations of ethanol. The resulting spectra showed two major peaks at 434 nm and 556 nm similar to what we observed with the deoxygenated form of the protein in absence of any ligand (**Fig. 5C**). These results imply that the heme group does not interact with ethanol. Rather, they suggest that ethanol binds the alpha helices of the HemAT sensing domain, perhaps using a mechanism similar to the one proposed for the McpB cytoplasmic signaling domain.

## Discussion

We found that *B. subtilis* performs chemotaxis to multiple short-chain alcohols. These alcohols are directly sensed by two chemoreceptors, McpB and HemAT. McpB is a transmembrane chemoreceptor, with an extracellular sensing domain and a cytoplasmic signaling domain, linked by a cytoplasmic HAMP domain. It is known to sense the amino acid asparagine and alkaline environments as attractants using the extracellular sensing domain (11, 24). HemAT, on the other hand, is a soluble chemoreceptor, which consists of a sensing and signaling domain but lacks a HAMP domain. Its myoglobin-like sensing domain contains heme and is known to bind molecular oxygen. Using chimeric receptors and STD-NMR, we found that short-chain alcohols are directly sensed by the cytoplasmic signaling domain of McpB and the sensing domain of HemAT. In the case of HemAT, the alcohols do not appear to bind heme; rather, they likely bind between the helices encapsulating the heme.

Among the alcohols tested, ethanol is the most likely physiological attractant, because it is produced by many microorganisms and is prevalent in nature (10). As a consequence, we focused on this chemical. Curiously, ethanol is not consumed by *B. subtilis*, suggesting that it is used for purposes other than nutrition. One possibility is that *B. subtilis* uses ethanol to locate prey, which could potentially explain why *B. subtilis* is attracted to a chemical that nominally inhibits its growth. The most likely prey are Crabtree-positive yeast such as *Saccharomyces cerevisiae*, which produce ethanol at high concentrations even during aerobic growth (37). Indeed, *B. subtilis* inhibits the growth of *S. cerevisiae* through the production of antifungal compounds (38) (**Fig. S6**).

In addition to fermenting microorganisms, plants can also produce ethanol at millimolar concentrations during anoxic conditions, such as when root systems are flooded with water (39). *B. subtilis* is known to form biofilm on root surfaces (40). These associations are symbiotic, where *B. subtilis* protects the plant from microbial pathogens and the plant provides nutrients to *B. subtilis*. Chemotaxis is known to promote colonization of plants roots by many species of bacteria (41). Typically, these bacteria are attracted to chemicals within root exudates. Indeed, one study found that *B. subtilis* is attracted by *Arabidopsis thaliana* root exudates (42). Therefore, another possibility is that *B. subtilis* also uses ethanol to locate roots for colonization.

Aside from ethanol, methanol may also be a physiological attractant. It is a byproduct of pectin degradation and, as a consequence, can contaminate alcoholic beverages (43). However, *B. subtilis* does not consume methanol because it lacks methanol dehydrogenase activity (44). Similar to ethanol, *B. subtilis* may use methanol to locate prey, this time pectin-degrading microorganisms.

A key difference between taxis to alcohols and conventional attractants, such as amino acids, is their respective sensitivities. Amino acids are sensed at micromolar concentrations whereas alcohols are sensed at millimolar concentrations. The weak affinity for alcohols is not surprising as most ethanol receptors in mammals also exhibit weak affinity for ethanol (45). The question then is whether ethanol is actually an attractant for *B. subtilis* given that relatively high concentrations are necessary to elicit taxis. Over-ripe fruits provide one potential source for high ethanol concentrations, where they can exceed one molar (46). In addition, plant roots can also provide another source at millimolar concentrations (47–49). This suggests that ethanol taxis can indeed occur in the environment. Whether the other alcohols reach such concentrations in the environment is not known.

While the physiological significance of alcohol taxis is still speculative at this stage, it nonetheless represents an unconventional attractant in the sense that it is not a nutrient per se but rather provides some other environmental cue. In this regard, it more closely resembles chemotaxis to homoserine lactones, involved in quorum sensing, or catecholamines, which are associated with gut colonization and virulence (50–52). The only other bacterium known to exhibit alcohol taxis is *Pseudomonas putida* (18). However, this bacterium consumes alcohols. In addition, it does not directly sense these alcohols but rather the byproducts of their degradation, namely carboxylic acid.

Perhaps the most interesting aspect of ethanol taxis involves the sensing mechanism. Typically, small-molecule attractants bind the extracellular sensing domain. The main exceptions are PTS sugars, which are sensed indirectly through the PTS system (5, 6). Ethanol is sensed intracellularly. In the case of HemAT, this distinction is minor, as ethanol binds the sensing domain, albeit one normally associated with oxygen sensing. In the case of McpB, the cytoplasmic signaling domain is involved in sensing ethanol through direct binding. This appears to be first documented case of the cytoplasmic signaling domain being directly involved in sensing. While we were able to establish that ethanol binds the McpB cytoplasmic signaling domain using genetics and STD-NMR, the detailed binding and induced signaling mechanisms are still somewhat opaque. In particular, it is not clear whether ethanol exerts its effect precisely at residue 431 or possibly at one or multiple other positions along the lengthy cytoplasmic domain. MD simulations suggest that ethanol can bind nonspecifically at several places on the McpB cytoplasmic surface as well as penetrate to the core of the four-helix bundle, at least within the methylation helix region. Although we did not observe ethanol to enter the bundle core near residue 431 in our simulations, it may do so on longer timescales or in particular signaling states. In addition, the precise molecular details of how ethanol binding induces signaling in wild-type McpB remain to be worked out. The enhanced packing interactions in the A431S McpB, which does not respond to ethanol, suggest that it may disrupt or loosen packing, leading to changes in the overall stability of McpB that can be transmitted to the kinase. This idea is in line with numerous previous studies of the *E. coli* Tsr and Tar chemoreceptors, for example, which suggest that changes in periplasmic ligand binding and adaptation state affect packing throughout the cytoplasmic bundle (5, 6).

Many aspects of ethanol sensing in *B. subtilis* are analogous to mechanisms observed in higher eukaryotes. In particular, alcohols generally bind proteins with low affinities, and relatively high concentrations of alcohols are required to induce behavioral effects. For example, receptors such as the N-methyl-D-aspartate-type glutamate receptors, γ-aminobutyric acid type A (GABA_A_) receptors, and glycine receptors all exhibit weak affinity for ethanol (> 10 mM) (45). Although the binding sites on these proteins are not well characterized, ethanol is thought to bind helical regions in most cases. In the case of GABA_A_ receptor, for example, it is thought that ethanol binds within a small cavity between two transmembrane helices (TM2 and TM3) (53). Similarly, in the case of odorant binding protein LUSH from *Drosophila melanogaster*, a small cavity between alpha helices accommodates a single ethanol molecule, where its hydroxyl group form hydrogen bonds with neighboring Thr^57^ and Ser^52^ residues (54).

Analysis of these binding sites suggests that ethanol preferentially binds helices with amphipathic surfaces (45, 55, 56). The sensing mechanisms for these proteins typically involve replacement of water molecules with ethanol within small hydrophobic cavities between two or more helices. Indeed, an analogous mechanism appears to be employed by the *B. subtilis* chemoreceptors.

## Materials and Methods

### Chemicals and growth media

The following media were used for cell growth: Luria-Bertani broth (LB: 1% tryptone, 0.5% yeast extract, and 0.5% NaCl); tryptone broth (TB: 1% tryptone and 0.5% NaCl); tryptose blood agar base (TBAB: 1% tryptone, 0.3% beef extract, 0.5% NaCl, and 1.5% agar); yeast-peptone-dextrose broth (YPD: 1% yeast extract, 2% peptone, and 2% dextrose); and capillary assay minimal medium (CAMM: 50 mM potassium phosphate buffer (pH 7.0), 1.2 mM MgCl_2_, 0.14 mM CaCl_2_, 1 mM (NH_4_)_2_SO_4_, 0.01 mM MnCl_2_, and 42 µM ferric citrate). Chemotaxis buffer consists of 10 mM potassium phosphate buffer (pH 7.0), 0.14 mM CaCl_2_, 0.3 mM (NH_4_)_2_SO_4_, 0.1 mM EDTA, 5 mM sodium lactate, and 0.05% (v/v) glycerol. All alcohols used in this study were purchased from Fisher Scientific, Inc.

### Strains and plasmids

All strains and plasmids used in this work are listed in Tables 1 and 2, respectively. Chemotaxis experiments were performed with derivatives of *B. subtilis* OI1085. Growth experiments were performed using *B. subtilis* 168, which is the parental strain. The undomesticated *B. subtilis* strain NCBI3610 and the *Saccharomyces cerevisiae* CEN.PK113-7D yeast strain were used in the antimicrobial diffusion assays. All cloning was performed using NEB® 5-alpha Competent *E. coli* (New England Biolabs). All oligonucleotides used in this study are provided in (**Table S2**).

**TABLE 2.** Plasmids used in this study.

| Plasmid | Description | Reference |
| --- | --- | --- |
| pET28a (+) | His-tagged cloning vector for protein purification; Kan<sup>R</sup> | Novagen |
| pJSpe | Modified pJOE8999 optimized for Gibson assembly of<br>homology templates; Amp<sup>R</sup>, Kan<sup>R</sup> | (24) |
| pPT037 | pJSpe::*cheV* (for *cheV* knockout) | (24) |
| pPT058 | pJSpe::*mcpB* (for *mcpB* knockout) | (24) |
| pPT053 | pJSpe::*hemAT* (for *hemAT* knockout) | This work |
| pAIN750 | *B. subtilis* empty vector for integration at *amyE*; Amp<sup>R</sup>,<br>Spc<sup>R</sup> | (83) |
| pPT200 | pAIN750::*mcpB*[M1-V287] *mcpA*[L287-E661] | This work |
| pPT205 | pAIN750::*mcpB*[M1-Q359] *mcpA*[D359-E661] | This work |
| pGB42 | pAIN750::*mcpB*[M1-A374] *mcpA*[S374-E661] | This work |
| pGB43 | pAIN750::*mcpB*[M1-N397] *mcpA*[E397-E661] | This work |
| pGB34 | pAIN750::*mcpB*[M1-I433] *mcpA*[Q433-E661] | This work |
| pGB64 | pAIN750::*mcpB*[M1-Q423] *mcpA*[A423-E661] | This work |
| pGB35 | pAIN750::*mcpB*[M1-L481] *mcpA*[R481-E661] | This work |
| pPT086 | pAIN750::*mcpA*[M1-Q358]-*mcpB*[D359-E662] | This work |
| pGB65 | pAIN750::*mcpB*[A431S] | This work |
| pGB83 | pAIN750::*mcpB*[T424A] | This work |
| pGB82 | pAIN750::*mcpB*[D427T] | This work |
| pGB67 | pAIN750::*mcpB*[E581Q] | This work |
| pGB79 | pAIN750::*mcpB*[K585E] | This work |
| pGB94 | pAIN750::*mcpB*[E399K] | This work |
| pPT262 | 6xHis-C terminal McpB expression plasmid,<br>pET28(a)::*mcpB<sub>C</sub>* | This work |
| pGB78 | 6xHis-C terminal McpB<sub>C</sub>[A431S] expression plasmid,<br>pET28(a)::*mcpB<sub>C</sub>*[A431S] | This work |
| pGEX-6p-2::*cheA* | GST-CheA overexpression plasmid | (15) |
| pGEX-6p-2::*cheW* | GST-CheW overexpression plasmid | (15) |
| pGEX-6p-2::*cheD* | GST-CheD overexpression plasmid | (15) |
| pGB46 | 6xHis-C terminal HemAT expression plasmid,<br>pET28(a)::*hemAT<sub>S</sub>* | This work |
| pSP03 | 6xHis-N terminal HemAT expression plasmid,<br>pET28(a)::*hemAT<sub>N</sub>* | This work |

Gene deletions were constructed using plasmids derived from pJSpe, which provides a CRISPR/Cas9-based, marker-free, and scarless genome editing system for *B. subtilis* (57). To construct a deletion vector, a 20-bp crRNA target sequence complementary to the targeted gene sequence was designed using the CHOPCHOP online tool (58). The 5’-end phosphorylated complementary oligonucleotides were then annealed and subcloned into BsaI restriction sites on pJSpe plasmid using Golden Gate assembly (59). The resultant plasmid was then linearized at SpeI restriction site and joined to two PCR fragments (∼700 to 800 bp) flanking the targeted gene using Gibson assembly (60). Prior to transformation into *B. subtilis* strain, each of the pJSpe-derived deletion plasmids was linearized at XhoI restriction site and subsequently self-ligated to create a long DNA concatemer. The concatemer was then transformed into *B. subtilis* strain using the two-step Spizizen method (61). Transformation product of *B. subtilis* strain and deletion plasmid concatemer was incubated on a LB agar (LB and 1.5% agar) plate supplemented with 5 µg/mL kanamycin and 0.2% mannose for about 24 h at 30 °C. Next, single colonies were isolated and twice streaked on fresh drug plates (described above) to assure a clonal genotype. Positive colonies were verified using colony PCR and again streaked on a plain LB agar plate and incubated for additional 24 h at 50 °C to cure the deletion plasmid. Colonies with cured plasmids were unable to grow on a LB agar plate supplemented with 5 µg/mL kanamycin.

To construct chemoreceptor chimeras, two opposing primers were designed to amplify DNA regions outward from the fusion points of the chimeric gene using PCR with pAIN750*mcpB* integration plasmid as the DNA template. Then, a second pair of primers with short overlapping regions were used to PCR amplify the desired fragment of *mcpA* gene from pAIN750*mcpA*. Following purification of PCR DNA products by gel extraction, the DNA fragments were assembled using Gibson assembly and transformed into *E. coli*. Following isolation from *E. coli* and sequence verification, the concatemer of the resultant integration plasmid was prepared as described above and transformed into *B. subtilis* OI3545, which lacks all ten chemoreceptors. Transformation product was then incubated on a LB agar plate supplemented with 100 µg/mL spectinomycin for 15 h at 37 °C. Single colonies were isolated and streaked on a TBAB agar (TBAB and 1.5% agar) plate supplemented with 1% soluble starch. A single positive colony with chemoreceptor expression cassette recombined to *amyE* locus was verified using Gram Iodine solution (0.33% iodine, 0.66% potassium iodide, and 1% sodium bicarbonate). Correct colonies with disrupted *amyE* gene were unable to form clear zones on TBAB-starch plate.

Point mutations on *mcpB* chemoreceptor gene were introduced using the inverse PCR method. Briefly, two opposing primers containing the desired mutations were used to PCR amplify integration pAIN750*mcpB* plasmid. Following purification of PCR DNA by gel extraction, 5’-ends of the DNA fragment was phosphorylated with T4 polynucleotide kinase and then blunt-end ligated using T4 DNA ligase. Ligation product was heat-inactivated and transformed into *E. coli*. Following isolation from *E. coli* and sequence verification, concatemer of the resultant integration plasmid was prepared as described above and transformed into *B. subtilis* OI3545 to integrate the mutant chemoreceptor expression cassette into the *amyE* locus.

Protein expression plasmids were constructed with the pET28(+) expression vector system using Gibson assembly. Briefly, DNA for the HemAT sensing domain (residues 1 to 178) was cloned in frame with a C-terminal His_6_-tag between the NcoI and HindIII restriction sites on pET28a(+). Similarly, the DNAs for the wild-type McpB and McpB[A431S] cytoplasmic regions including the HAMP domain (residues 305 to 662) were cloned in frame with a C-terminal His_6_ tag at NcoI restriction site on pET28a(+). The DNA for HemAT signaling domain (residues 177 to 432) was cloned in frame with a N-terminal His_6_-tag at the NheI restriction site on pET28a(+). After isolation and sequence verification, all plasmids were transformed into *E. coli* BL21 (DE3) strain for protein expression and purification.

### Protein expression and purification

CheA, CheW, and CheD proteins used in the kinase assay were expressed from glutathione *S*-transferase (GST) fusion plasmids and purified from *E. coli* BL21(DE3) strain as described previously (15, 23). GSTrap columns (5 mL; GE Healthcare) were used with an Akta Prime FPLC system (GE Healthcare) for purification. To purify the GST fusion proteins, cells were grown in 2 liters of LB with 100 μg/mL ampicillin at 37 °C and shaking at 250 rpm until OD_600_ = 0.8. Expression was then induced by the addition of 1 mM IPTG (isopropyl-β-d-thiogalactopyranoside), and the culture was grown for 12 h at 25 °C with 250 rpm shaking. For CheA, the culture was induced at 37 °C for 4 h. Cells were then centrifugated at 8000 x *g* for 8 min and resuspended in Tris-buffered saline (TBS: 50 mM Tris, 150 mM NaCl, pH 7.5) supplemented with 1% Triton X100 and 1 mM of dithiothreitol (DTT) for every 1 g of cell pellet. The cells were then disrupted by sonication (5 x 10 s pulse). The supernatants were clarified by two rounds of centrifugations (9,000 x *g*, 15 min; 40,000 x *g*, 40 min), and loaded onto 5 mL GSTrap columns pre-washed with 10 column-volumes of TBS. Protein-bound columns were then washed with at least 15 volumes of TBS, and GST tagged proteins were eluted using 10 mL glutathione elution buffer (GEB: 50 mM Tris, 5 mM glutathione, pH 8). To remove the GST tag, the purified proteins were cleaved by PreScission protease, as specified by the supplier (Amersham Biosciences), and applied to another 5 mL GSTrap column. The flow-through was collected and concentrated to approximately 5 mL using a cellulose ultrafiltration membrane (Millipore) in an Amicon ultrafiltration cell. Last, the purified proteins were dialysed in TKMD buffer (50 mM Tris, 50 mM KCl, 5 mM MgCl_2_, 0.1 mM DTT, pH 8) and aliquots were stored at −80°C.

*E. coli* BL21 (DE3) cells harboring the His_6_-tagged expression plasmids were grown in 2 L of LB medium supplemented with 30 µg/mL kanamycin at 37 °C and shaking at 250 rpm until *A*_600_ = 0.7. Expression was then induced by the addition of 1 mM IPTG, and the cultures were grown for 12 h at 25 °C. Cells were harvested by centrifugation at 7,000 x *g* at 4°C for 10 min. Cells harboring HemAT_N_ were resuspended in lysis buffer (50 mM NaH_2_PO_4_, 300 mM NaCl, 10 mM Imidazole, pH 8) and sonicated (5 x 10 s pulses). Cell debris was removed by centrifugation at 12,000 x *g* for 1 h. The dark-red supernatant containing HemAT_N_ was loaded on a 5 mL GE HisTrap column prewashed with NiSO_4_ and binding buffer (50 mM NaH_2_PO_4_, 300 mM NaCl, 20 mM imidazole, pH 8). The protein-bound column was then washed with binding buffer and proteins were eluted with elution buffer (50 mM NaH_2_PO_4_, 300 mM NaCl, 250 mM Imidazole, pH 8). The collected HemAT_N_ protein samples were concentrated using an Amicon ultrafiltration cell (Millipore) and dialyzed into dialysis buffer (50 mM Tris, 300 mM NaCl, pH 8) at 4 °C and aliquots were stored at −80°C.

McpB_C_, McpB_C_ [A431S], and HemAT_S_ proteins were purified under denaturing conditions. Briefly, cells were induced and grown as described above. Cells were then resuspended in buffer B (8 M urea, 0.1 M NaH_2_PO_4_, 0.01 M Tris, pH 8) with 1% Triton X100 and 1 mM of DTT for every 1 g of cell pellet and incubated at room temperature for 1 h. Cell suspension was clarified by centrifugation at 40,000 x *g* for 1 h. The cell lysates were loaded onto 5 mL GE Hi-Trap Chelating column charged with 0.1 M NiSO_4_ and washed with buffer B and buffer C (buffer B at pH 6.3). The fusion proteins were eluted from the column with 25 mL elution buffer E (buffer B at pH 4.5). Proteins were refolded by dialyzing in PBS (10 mM Na_2_HPO_4_, 1.8 mM KH_2_PO_4_, 137 mM NaCl, 2.7 mM KCl, pH 7.4) at 4 °C, and aliquots were stored at −80 °C. Purified proteins proper folding was verified with circular dichroism spectroscopy. Concentration of all purified proteins were quantified by Pierce BCA protein assay kit. SDS-PAGE images of the purified recombinant chemoreceptor proteins are shown in **Fig. S7**.

### Capillary assay for chemotaxis

The capillary assay was performed as described previously (62). Briefly, cells were grown for 16 h at 30 °C on TBAB plates. The cells were then scraped from the plates and resuspended to OD_600_ = 0.03 in 5-mL CAMM supplemented with 50 μg/mL histidine, 50 μg/mL methionine, 50 μg/mL tryptophan, and 20 mM sorbitol, and 2% TB. The cultures were grown to OD_600_ = 0.4 – 0.45 at 37 °C with shaking at 250 rpm. At this point, 50 μL of GL solution (5% (v/v) glycerol and 0.5 M sodium lactate) was added, and cells were incubated for another 15 min (at 37 °C and 250 rpm shaking). The cells were then washed twice with chemotaxis buffer and incubated for additional 25 min (at 37 °C and 250 rpm shaking) to assure that the cells were motile. Cells were then diluted to OD_600_ = 0.001 in chemotaxis buffer and aliquoted into 0.3-mL ponds on a slide warmer at 37 °C and closed-end capillary tubes filled with alcohol solutions or asparagine solution (3.16 μM) prepared in the same chemotaxis buffer were inserted. After 30 min, cells in the capillaries were harvested and transferred to 3 mL of top agar (1% tryptone, 0.8% NaCl, 0.8% agar, and 0.5 mM EDTA) and plated onto TB agar (TB and 1.5% agar) plates. These plates were incubated for 16 h at 37 °C and colonies were counted. Experiments were performed in triplicate each day and repeated on three different days.

### Cell growth

Cells density was measured as optical absorbance at 600 nm. Briefly, *B. subtilis* 168 was first grown for 16 h at 30°C on a TBAB plate. For growth experiments in minimal medium, the cells were first scraped from the TBAB plate and then resuspended to OD_600_ = 0.03 in 50 mL CAMM supplemented with 50 μg/mL tryptophan and 5 g/L glucose; and grown at 37 °C with shaking at 250 rpm. At the OD_600_ = 0.8, the cells were diluted 1:20 (v/v) into 50 mL CAMM containing 50 μg/mL tryptophan, supplemented with 0.01 M ethanol, 0.1 M ethanol, or 5 g/L glucose (positive control), respectively, and grown for 24 h at 37 °C with shaking at 250 rpm. For growth experiments in rich medium, cell cultures starting at OD_600_=0.03 were grown to OD_600_ = 0.4 at 37°C with shaking at 250 rpm in 50 mL LB media. At this point, cell cultures were supplemented with 0.01 M, 0.1 M, or 1.0 M ethanol, respectively, and grown for another 5 h at 37°C with shaking at 250 rpm. All growth experiments were performed in triplicate.

### Ethanol utilization experiments

Ethanol concentrations were measured using a Shimadzu high-performance liquid chromatography system equipped with a RID-10A refractive index detector, an Aminex HPX-87H carbohydrate analysis column (Bio-Rad Laboratories), and a cation H microguard cartridge (Bio-Rad Laboratories). The column and guard cartridge were kept at 65 °C, and 0.5 mM H_2_SO_4_ was used a mobile phase at a constant flow rate of 0.6 mL/min. Prior to measurements, cells in culture samples were pelleted, and the resulting supernatant was passed through a 0.22-µm polyethersulfone syringe filter. Peaks were identified and quantified by retention time comparison to the standards.

### Alcohol dehydrogenase activity measurement

*B. subtilis* OI1085 was first grown for 16 h at 30 °C on a TBAB plate. For aerobic growth, the cells were then scraped from the TBAB plate and resuspended to OD_600_ = 0.03 in 5 mL CAMM supplemented with 50 µg/mL histidine, methionine, tryptophan, 20 mM sorbitol, and 2% TB, and grown at 37 °C with vigorous shaking at 250 rpm. For anaerobic growth, however, cells were cultured starting at OD_600_ = 0.03 in a sealed bottle filled to the top without agitation in CAMM supplemented with 1% glucose and mixture of all 20 amino acids at 50 µg/mL (20). For *E. coli* cultures, the cells (MG1655) were grown in M9 media supplemented with 0.4% glucose at 37°C in sealed bottles filled to the top without agitation for anaerobic growth and in flasks with shaking at 250 rpm for aerobic growth (22). All cell cultures were grown to stationary phase prior to sonication (7 x 10 s pulses), and soluble cell extracts were obtained by centrifugation (7000 x *g* at 4°C for 10 min). Alcohol dehydrogenase enzyme assays were performed as described previously (63). Briefly, the assay reactions were prepared with 22 mM sodium pyrophosphate (pH 8.8), 0.3 mM sodium phosphate, 7.5 mM β-nicotinamide adenine dinucleotide, 0.003% (w/v) bovine serum albumin, 1.6% (v/v) of desired cell lysate, and 3.2% (v/v) ethanol in 200 µL reaction volume. Then, the reduction of NAD^+^ to NADH was recorded at 340 nm using a Shimadzu UV-1800 spectrophotometer. One unit of alcohol dehydrogenase activity is defined as the amount of enzyme that converts 1 µmole of ethanol to acetaldehyde per minute at pH 8.8 at 25 °C.

### Antimicrobial diffusion assay

Antifungal activity of the *B. subtilis* strains were assayed using the disc diffusion method as described previously (64). Briefly, the *S. cerevisiae* CEN.PK113-7D was grown in YPD rich medium for 24 h at 30 °C with shaking at 200 rpm. 0.1% (v/v) of yeast culture was mixed with YPD top agar (YPD with 0.8% agar) and spread on top of a YPD plate (YPD with 2% agar). Once top yeast layer was solidified, 10-mm filter paper (Whatman Filter Paper, Grade 1) discs loaded with supernatants from *B. subtilis* strains grown overnight in LB medium at 37°C, were placed on top of the yeast layer. As negative controls, separate discs were loaded with LB and water. The plate was incubated at 30°C for another 24 h, and then imaged. A zone of inhibition around the discs indicated antifungal activity.

### Preparation of bacterial membranes

Cells were grown for 16 h at 30 °C on TBAB plates. The cells were then scraped from the plates and resuspended to OD_600_ = 0.03 in 50-mL CAMM supplemented with 50 μg/mL histidine, 50 μg/mL methionine, 50 μg/mL tryptophan, 20 mM sorbitol, and 2% TB. The cells were grown at 37 °C with aeration until they reach mid-exponential phase. The cells were then diluted 1:10 (v/v) into 50 mL CAMM media and grown till mid-exponential phase. The cells were again diluted to an OD_600_ of 0.01 in 50 mL media and grown till mid-exponential phase. Finally, the cultures were diluted 1:10 (v/v) into multiple flasks containing 50 mL media and grown with shaking at 37 °C until an OD_600_ of 0.6. The cells were then harvested by centrifugation at 9900 x *g* for 15 min and washed 3 times with 1 M KCl to remove extracellular proteases. Cells were resuspended in sonication buffer+ (10 mM potassium phosphate (pH 7), 10 mM MgCl_2_, 1 mM EDTA, 0.3 mM DTT, 20 mM KCl, 1 mM glutamate, 2 mM phenylmethanesulphonyl fluoride, and 20% glycerol). EDTA and phenylmethanesulphonyl fluoride were added as protease inhibitors. Cells were sonicated, and the cell debris was removed by centrifugation at 17,600 x *g* at 4 °C for 15 min. Bacterial membranes were removed by centrifugation at 120,000 x *g* for 2 h at 4 °C in a Beckman 70 Ti rotor. Pelleted membranes were resuspended in MT buffer (10 mM potassium phosphate (pH 7), 1 mM MgCl_2_, 0.1 mM EDTA, and 1 mM 2-mercaptoethanol), and homogenized using a glass/Teflon homogenizer followed by another centrifugation at 120,000 x *g* for 2 h at 4 °C. This step was repeated once more. Finally, the membranes were homogenized in MT buffer at a concentration of 32 mg/mL and stored in small aliquots at −80 °C.

### *In vitro* assay for receptor-coupled kinase activity

Reactions consisted of purified *B. subtilis* membranes expressing McpB or HemAT as the sole chemoreceptor and purified CheW, CheA, and CheD prepared in buffer (50 mM Tris, 50 mM KCl, 5 mM MgCl_2_, pH 7.5) at the following concentrations: 6 µM chemoreceptor, 2 µM CheW, 2 µM CheA kinase, and 2 µM CheD. Ethanol was then added to the mixture at different final concentrations in 20 µL reaction volume. As a negative control, only buffer was added. Reactions were then pre-incubated at 23 °C for 1 h to permit the formation of the chemoreceptor-kinase complex. CheA autophosphorylation was initiated by the addition of [γ-^32^P] ATP (4000-8000 cpm/pmol) to a final concentration of 0.1 mM. 5 µL aliquots were quenched at 15 s by mixing the reactions with 15 µL of 2X Laemmli sample buffer containing 25 mM EDTA at room temperature, essentially fixing the level of phosphor-CheA. Initial phosphor-CheA formation rates were analyzed using 12% SDS-PAGE. Gels were dried immediately after electrophoresis and phosphor-CheA was quantified by phosphor-imaging (Molecular Dynamics) and ImageJ (65).

### Circular dichroism (CD) spectroscopy

CD-spectra was measured on a JASCO J-720 spectropolarimeter (Japan Spectroscopic Co., Inc., Tokyo, Japan) with a cuvette of path length 0.1 cm. Prior to measurements, protein samples were dialyzed into 10 mM sodium phosphate buffer (pH 8) and diluted to 5 µM. Spectral measurements were carried out in triplicate using a scanning rate of 50 nm/min. A buffer only control sample was used for baseline correction. Structural analysis was done using BeStSel (66).

### Ultraviolet-visible (UV) spectral measurements

All UV-spectral measurements were performed on a Shimadzu UV-1800 spectrophotometer. The UV-spectra of the oxygenated sensing domain of HemAT (HemAT_N_) protein was measured in aerobic conditions. To measure the UV-spectra of HemAT_N_ in presence of ethanol, protein samples were first deoxygenated by adding a few grains of sodium dithionite in a glove box. Sodium dithionite-reduced protein samples were then titrated with different doses of ethanol in sealed quartz cuvettes, and the UV-spectra (200 nm to 600 nm) of these samples were immediately recorded in the spectrophotometer.

### Saturation-transfer difference nuclear magnetic resonance spectroscopy (STD-NMR)

All NMR spectroscopy measurements were performed on a Varian VNMRS instrument at 750 MHz with 5 mm Varian HCN probe at 298 K without sample spinning. Prior to measurements, protein samples were buffer exchanged into PBS (50 mM KH_2_PO_4_, 20 mM NaCl, pH 7.4) in D_2_O using Micro Bio-Spin® Columns with Bio-Gel® P-6 (Bio-Rad Laboratories, Hercules, CA, USA). To avoid aggregation, HemAT_N_ protein was buffer exchanged into modified PBS (50 mM KH_2_PO_4_, 300 mM NaCl, pH 8.0) containing 10% D_2_O. 50 µM Protein samples were then mixed with ethanol (final concentration of 3 mM) in a 500 µL solution. ^1^H spectra were obtained from 32 scans with a 90-degree pulse and a 2-s relaxation delay. In STD-NMR experiments, the protein samples were selectively saturated at 2.15 ppm with a train of Gaussian pulses of 50 ms duration with 0.1 ms delay and 5 s relaxation delay for a total saturation time of 3 s and 2048 scans. Off-resonance irradiation was applied at 30 ppm. Trim pulse of 50 ms was used to reduce protein background. In the case of HemAT_N_, the protein sample was saturated at 7.06 ppm and 256 scans were used to obtain spectra. All STD spectra were obtained by internal subtraction via phase cycling after a block size of 8 to reduce artifacts resulted from temperature variation and magnet instability. Control experiments were performed on samples containing only ethanol without protein. All areas were calculated using MNova V14.1 (by Mestrelab chemistry solutions) in stacked mode.

### Structural analysis

Domains of the McpB, McpA, TlpA, and TlpB chemoreceptors from *B. subtilis* were predicted using phmmer search engine on the HMMER web server using the UniProt reference proteomes database with default sequence E value thresholds (67). The amino acid sequences of the cytoplasmic signaling domains were then manually obtained based on the previous large-scale alignment results (9). To identify the three structural subdomains of the cytoplasmic signaling domain, the sequences were then aligned with the amino acid sequences of the corresponding domains from the Tar, Tsr, Trg, and Tap chemoreceptors of *E. coli* using MUSCLE (68) with the default parameter values. Pairwise amino-acid sequence alignments between the protein-pairs (McpA-McpB; McpA-HemAT; and HemAT-YfmS) for chimeric receptor analysis were performed using EMBOSS Water (69). A homology model of the cytoplasmic signaling domain of the McpB dimer (residues 352 to 662) was constructed in Modeller (v-9.23) (70) using the *Thermatoga maritima* Tm113 chemoreceptor (PDB 2CH7) as the template (71). Side chain conformations were refined using SCWRL4 (72) and the entire structural model was subsequently refined using the YASARA energy minimization server (73). The resulting Ramachandran plots were verified using Procheck (74). The crystal structure of HemAT sensing domain from *B. subtilis* (PDB 1OR6) (33) was used for visualization. Visualization of all structures was accomplished using the VMD software package (v-1.9.3) (75).

### Receptor-ligand *in silico* docking experiment

The putative binding sites for ethanol were determined using Autodock (v-4.0) (76). Briefly, hydrogen atoms were first added to the McpB cytoplasmic signaling domain dimer model, and the number of torsional degrees of freedom for ethanol were set at 1. Autogrid was then used to adjust the position of grid boxes (60 x 60 x 60 points with 0.375 Å spacing for each box) on the ethanol-sensing region (residues 390 to 435). Finally, the Lamarckian genetic algorithm was employed to obtain the best docking site configurations.

### Molecular dynamics simulations

All-atom molecular dynamics simulations were conducted using NAMD 2.13 (77) and the CHARMM36 force field (78). Simulations were carried out in the NPT ensemble (pressure = 1 atm, temperature = 310 K) with values for general simulation parameters as previously described (79). The McpB cytoplasmic dimer model was solvated with TIP3P water and 150 mM NaCl using VMD (75), and 165 ethanol molecules (0.316 M) were randomly placed within the simulation box using the *gmx insert-molecules* tool. A copy of the system that included the A431S mutation was created, and both the wild-type and mutant McpB/ethanol systems were subjected to a conjugant-gradient energy minimization (2,000 steps) followed by a 10 ns equilibration simulation with protein backbone restraints and 3 x 600 ns unrestrained production simulations.

### MD simulation analysis

Density maps representing the average ethanol occupancy were computed using the VolMap plugin in VMD with default settings and averaging over each production simulation for the wild-type and A431S McpB/ethanol systems. To highlight unique binding sites between the two maps, a difference map was computed by subtracting the A431S map from the wild-type using VMD’s volutil plugin and removing smaller volumes resulting from slight irregularities in overlapping sites using the ‘hide dust’ feature in UCSF Chimera. All densities are visualized at an isovalue of 0.03 besides the difference map, which used an isovalue of 0.015. Protein-ethanol coordination was computed by measuring the minimum distance between non-hydrogen atoms in each residue and the nearest ethanol molecule; if this distance was less than 4 Å, the pair was considered to be in contact. Average coordination values were computed for each residue in the wild-type and A431S mutant McpB/ethanol systems by averaging over all three production simulations at 200 picosecond intervals. Percent changes were obtained by subtracting the values obtained in the latter from the former. Knobs-in-holes packing within the McpB cytoplasmic signaling domain was analyzed using the program SOCKET (80) with a packing cutoff of 7.8 Å (81). For each production simulation, knobs-in-holes packing was assessed at 2-ns intervals over the course of the trajectory, not including the first 100 ns to allow for packing changes resulting from equilibration or the A431S mutation. The occupancy of a particular knob-in-hole interaction over a given simulation was taken as the number of intervals in which it was identified by SOCKET divided by the total number of intervals analyzed in the simulation. The reported knobs-in-holes occupancies were averaged over both McpB monomers and all three production simulations for each McpB/ethanol system; error bars denote one standard deviation from the mean.

## Data availability

Raw data for all experiments are provided as Data set S1.

## Acknowledgment

We thank Prof. Jodi A. Hadden-Perilla for discussions surrounding the use of SOCKET. We would also like to thank Dr. Lingyang Zhu, Dr. Dean Olson, and the SCS NMR laboratory at the University of Illinois at Urbana-Champaign for valuable inputs and help with NMR measurements; Dr. Ahmed Hetta and Dr. Issac Caan for help with CD Spectroscopy experiments. This work was partially funded by National Institutes of Health Grant GM054365 and by the University of Illinois through the Robert W. Schaefer Faculty Scholar fund.

## Supplementary Information

**TABLE S1.** Putative ethanol-binding sites within the McpB ethanol-sensing region predicted by *in silico* docking experiments.

**TABLE S2.** Oligonucleotides used in this study

**FIG S1. Ethanol induces receptor-coupled kinase activity.** Levels of phosphorylated CheA kinase protein complexed with CheW, CheD, and (A) McpB or (B) HemAT chemoreceptors within the isolated membranes were measured in presence of increasing ethanol concentrations. 3.16 µM asparagine was used as positive control for membranes containing McpB, and buffer was used as negative control in all experiments.

**FIG S2. Amino-acid sequences of three structural subdomains within the cytoplasmic signaling domains of *B. subtilis* transmembrane chemoreceptors.** Amino-acid sequences of three structural subdomains, known as methylation helix (MH), flexible bundle (FB), and conserved signaling (CS)), within the cytoplasmic signaling domains of four *B. subtilis* transmembrane chemoreceptors are shown. For comparison, aligned amino-acid sequences of the corresponding subdomains from four *E. coli* transmembrane chemoreceptors are also shown. Characteristic seven-residue repeats (heptads) along the helices are labeled *a* to *g* and the corresponding amino-acid sequences are separated by alternating gray and white colors.

**FIG S3. Identification of putative ethanol-binding sites on the cytoplasmic signaling domain of McpB dimer.** (A) Five different clusters of putative binding sites within the ethanol sensing region spanning residues (390 to 435) on the N-helix and neighboring residues (577 to 622) on the C-helix of the McpB dimer fragment, predicted by *in silico* docking experiments. Monomers A and B are shown in blue and green, respectively. (B) Amino-acid sequence alignment of the ethanol-sensing region spanning residues (392 to 434) on the N-helix and neighboring residues (578 to 620) on the C-helix of McpB and the corresponding regions on McpA, TlpA, and TlpB. Conserved and non-conserved putative ethanol-binding residues are highlighted in red and green, respectively.

**FIG S4. CD-spectra of purified recombinant proteins.** The CD-spectra of recombinant wild-type and mutant (A431S) McpB cytoplasmic regions spanning residues (305 to 662), and recombinant C-terminal HemAT signaling domain spanning residues (177 to 432).

**FIG S5. Ethanol and knob-residues occupancy along the McpB coiled-coil.** (A) Density maps of the average ethanol occupancy along the wild-type (purple) and the A431S mutant (orange) McpB cytoplasmic signaling domain (McpB_C_). Differences between the wild-type and the A431S mutant (red density) reveal three distinct putative ethanol binding sites (S1, S2, and S3). Average changes in protein-ethanol coordination highlight the putative amino-acid residues (red bars) involved in ethanol binding in each site. (B) Distribution of knob residues (purple, space-filling) on the McpB cytoplasmic signaling dimer as identified using SOCKET (hole residues are not shown). The close-up depicts the identified knobs nearby the residue Ala^431^ (yellow). In addition to Ala^431^, residues Ala^583^ and Lys^585^ (cyan) are predicted to have higher average knob occupancies in McpB_C_-A431S compared to the wild-type McpB_C_. Data and error bars associated with the knob occupancies in panel b denote the means ± standard deviations from three independent simulations.

**FIG S6. Antifungal activity of *B. subtilis* strains.** (A) Growth inhibition of *S. cerevisiae* by supernatants from overnight cell cultures of *B. subtilis* OI1085 laboratory chemotaxis strain and undomesticated NCBI 3610 strain is measured using disk diffusion assay. Similar experiments were conducted using only water or LB instead of culture supernatant, as negative controls. (B) Chemotaxis responses of *B. subtilis* OI1085 laboratory chemotaxis strain and undomesticated NCBI 3610 strain to 1.78 M ethanol and buffer. Data and error bars shown in panel b denote the means ± standard deviations from three biological replicates performed on at three separate days.

**FIG S7.** SDS-PAGE images of purified recombinant chemoreceptor proteins used in this study.

**DATA SET S1.** Raw data for all experiments reported in the manuscript.

